# Bottom-up sensory processing can decrease activity and functional connectivity in the default mode like network in rats

**DOI:** 10.1101/482638

**Authors:** Rukun Hinz, Lore M. B. Peeters, Disha Shah, Stephan Missault, Michaël Belloy, Verdi Vanreusel, Meriam Malekzadeh, Marleen Verhoye, Annemie Van der Linden, Georgios A. Keliris

## Abstract

The default mode network is a large-scale brain network that is active during rest and internally focused states and deactivates as well as desynchronizes during externally oriented (top-down) attention demanding cognitive tasks. However, it is not sufficiently understood if unpredicted salient stimuli, able to trigger bottom-up attentional processes, could also result in similar reduction of activity and functional connectivity in the DMN. In this study, we investigated whether bottom-up sensory processing could influence the default mode like network (DMLN) in rats. DMLN activity was examined using block-design visual functional magnetic resonance imaging (fMRI) while its synchronization was investigated by comparing functional connectivity during a resting versus a continuously stimulated brain state by unpredicted light flashes. We demonstrated that activity in DMLN regions was decreased during visual stimulus blocks and increased during blanks. Furthermore, decreased inter-network functional connectivity between the DMLN and visual networks as well as decreased intra-network functional connectivity within the DMLN was observed during the continuous visual stimulation. These results suggest that triggering of bottom-up attention mechanisms in anesthetized rats can lead to a cascade similar to top-down orienting of attention in humans and is able to deactivate and desynchronize the DMLN.

## 1. Introduction

The brain is a complex network consisting of functionally interconnected regions that dynamically communicate with each other. Part of these interactions can be observed non-invasively using resting-state functional magnetic resonance imaging (rsfMRI) (Damoiseaux et al., 2006; Salvador et al., 2005; van den Heuvel and Hulshoff Pol, 2010). This technique relies on the detection of low frequency fluctuations (0.01-0.2 Hz) in the blood oxygen level dependent (BOLD) signal while the subject is at rest, *i.e.* not performing any task. The coordinated fluctuations in the signals of anatomically separated regions have been shown to reflect intrinsic brain networks and evidence suggests that they correspond to spontaneous neuronal activity (Krishnan et al., 2018; Ma et al., 2016; Petridou et al., 2013). The regions that show temporally highly correlated activity are considered to be functionally connected and are referred to as resting state networks (RSNs) (Friston, 2011).

Since its discovery, rsfMRI has been widely used in human research to study RSNs in the healthy brain as well as their alterations in neuropathologies (Greicius, 2008; Hull et al., 2017; Zhou et al., 2017). More recently, comparable RSNs have also been detected in rodents (Gozzi and Schwarz, 2016; Jonckers et al., 2011; C.P. Pawela et al., 2008; Sierakowiak et al., 2015). This finding has been very important as it opened a new window of pre-clinical investigations in (genetic) animal models of disease that can be investigated with different modalities at multiple scales, providing additional information about the underlying mechanisms (Nestler and Hyman, 2010; Trancikova et al., 2011). In addition, pre-clinical rsfMRI shows great potential in identifying early biomarkers for multiple neuropathologies and can be used as an excellent theranostic tool (Bertero et al., 2018; Li et al., 2017; Shah et al., 2016, 2013). However, the translation and interpretation of RSNs between rodents and humans remains challenging, among other reasons, due to the differences in anatomy, physiology and the required use of anesthesia in rodents (Pan et al., 2015). It is therefore of utmost importance to investigate and improve our understanding of specific rodent RSNs that have been suggested to be homologous to RSNs in humans.

A RSN that has raised a lot of interest in humans is the default mode network (DMN), which has been shown to be most active during rest and internally focused tasks and less active during externally oriented attention demanding cognitive tasks (Greicius et al., 2003). Thus, it has been classified as a “task-negative network” and has been shown to alternate its activity with “task-positive networks” (Fox et al., 2005). The DMN is thought to play a fundamental role in self-referential thought, mind-wandering, internally-oriented cognition, and autobiographical memory (Lin et al., 2017). In humans, this network consists of regions in the anterior pre-frontal cortex, posterior cingulate cortex/retrosplenial cortex (precuneus), hippocampal formation, medial and lateral parietal regions (Buckner et al., 2008; Laird et al., 2009; Liska et al., 2015). A default mode like network (DMLN) suggested to be homologous to the human DMN has also been identified in rodents (Lu et al., 2012; Stafford et al., 2014). This DMLN comprises comparable regions, *i.e.* orbital cortex, prelimbic cortex, cingulate cortex, temporal association cortex, auditory cortex, posterior parietal cortex and parietal association cortex, retrosplenial cortex and hippocampus. Furthermore, besides anatomical similarities, few studies could indicate functional similarities such as the higher activity of the DMLN during rest vs task and the anti-correlation relationship of the DMLN with the task positive network (Rohleder et al., 2016; Schwarz et al., 2013).

In recent years, multiple human studies have shown that functional connectivity (FC) within the DMN is decreased when subjects are in a higher attentive and cognitive brain state associated with performing an internally guided (top-down) attention-demanding task (Elton and Gao, 2015; Fransson, 2006; Gao et al., 2013; Marrelec and Fransson, 2011). It is thought that this internally guided attention to external sensory input can suppress other internal processes associated with the DMN, resulting in this network’s inactivation and relative disconnection (Gao et al., 2013; Mayer et al., 2010). However, it is not yet sufficiently understood if attentional guidance by externally driven factors (bottom-up) could also result in similar reduction of activity and connectivity in the DMN. In humans, some studies suggested that the DMN is not deactivated by simple sensory processing (Greicius et al., 2003). It should be noted, however, that the stimulus design in these studies was simple and predictable and thus not expected to continuously drive bottom-up attention. Interestingly, a study using simple but unpredictable visual stimuli could dynamically activate attention network and DMN indicating their interaction during stimulus-driven processes of attention (Hahn et al., 2007).

Neurophysiological experiments in the past few years suggest that top-down and bottom-up processes share overlapping neural systems and in particular the employment of the prefrontal and parietal network (for a review see (Katsuki and Constantinidis, 2014)). We conjectured, that similarly to top-down, bottom-up attention triggering stimuli could also deactivate DMN and reduce its connectivity. To test this hypothesis, we performed fMRI experiments in anesthetized rats driven by randomized (unpredictable) continuous visual stimulation (CVS) and compared DMLN activity and connectivity with a resting state baseline scan (RSB) and a blocked visual stimulation (BVS) design.

## 2. Material and Methods

### 2.1. Animals and ethical statement

In this study, we used male Long Evans wild type rats (N=12) of 4 months of age (Long Evans, Janvier). Rats were kept under a normal day/night cycle (12/12) with an average room temperature of 20-24°C and 40% humidity. Furthermore, rats were group housed and had *ad libitum* access to standard rodent chow and water. One animal was excluded from the analysis due to the detection of unilateral ventricular enlargement. All procedures were performed in accordance with the European Directive 2010/63/EU on the protection of animals used for scientific purposes. The protocols were approved by the Committee on Animal Care and Use at the University of Antwerp, Belgium (permit number: 2015-50), and all efforts were made to minimize animal suffering.

### 2.2. Animal preparation and anesthesia

Rats were first anesthetized using 5% isoflurane for induction and 2% isoflurane for maintenance (IsoFlo, Abbott Illinois, USA) in a mixture of 70% N_2_ and 30% O_2_. Animals were head fixed in the scanner using bite- and ear-bars and ophthalmic ointment was applied to the eyes. As soon as the animal was fixed in the scanner, medetomidine anesthesia (Domitor, Pfizer, Karlsruhe, Germany) was administered via a subcutaneous bolus of 0.05 mg/kg and isoflurane concentration was decreased to 0% over a time period of 5 minutes. Continuous subcutaneous infusion of medetomidine anesthesia of 0.1 mg/kg.h was started 15 minutes post bolus injection. Functional MRI scans were acquired starting from 30 min post-bolus injection until 1h05 min post bolus injection. The physiological status of the animals was monitored throughout the entire imaging procedure. Respiratory rate was obtained from a pressure sensitive pad (MR-compatible Small Animal Monitoring and Gating system, SA Instruments, Inc.) with a sampling rate of 225 Hz. Body temperature was closely monitored using a rectal thermistor and was maintained between (37.0 ± 0.1 °C) using a feed-back controlled warm air heat system (MR-compatible Small Animal Heating System, SA Instruments, Inc.). Furthermore, blood oxygenation was recorded using a pulse oxygenation meter (MR-compatible Small Animal Monitoring and Gating system, SA Instruments, Inc.) with a sampling rate of 450 Hz. After imaging procedures, animals received a subcutaneous bolus injection of 0.1 mg/kg atipamezole (Antisedan, Pfizer, Karlsruhe, Germany) to counteract the effects of medetomidine and were placed in a recovery box with infrared heating for post-scan monitoring until the animal was fully awake.

### 2.3. MRI

All imaging procedures were performed with Paravision 6.0 using a 9.4 T BioSpec MR system (Bruker, Germany) with an active decoupled rat quadrature surface coil (Rapid biomedical, Germany) and a 98 mm diameter quadrature volume resonator for transmission (Bruker, Germany). First, three orthogonal anatomical multi-slice Turbo RARE T2-weighted images (field of view (FOV): (30×30) mm^2^, matrix dimensions (MD): [256×256], 12 slices, Slice thickness (ST): 0.9 mm, echo time (TE)/ repetition time (TR): 33/2500 ms, RARE factor: 8) were acquired to allow reproducible flat skull positioning of coronal slices. Then, a coronal anatomical reference scan was acquired using a multi-slice Turbo RARE T2-weighted sequence (FOV: (30×30) mm^2^, MD: [256×256], 12 slices, ST: 0.9 mm, TE/TR: 33/2500 ms, RARE factor: 8), covering the brain from 3.3 mm anterior to bregma to 7.5 mm posterior to bregma (suppl. figure 1). Next, a B_0_ field map was acquired to assess field homogeneity, followed by local shimming in a rectangular volume of interest in the brain to correct for the measured inhomogeneities. The RSB scan had an identical geometry to the reference scan and was acquired using a T2*-weighted single shot echo planar imaging (EPI) sequence (FOV: (30×30) mm^2^, MD: [128×98], 12 slices, ST: 0.9 mm, TE/TR: 18/2000 ms) resulting in a voxel dimension of (0.234 × 0.313 × 0.9) mm^3^ and a total scan duration of 10 min (300 volumes). Subsequently, random continuous light flickering (see section 2.4) was turned on and after 1 minute the CVS scan was acquired using the same parameters as the RSB (300 volumes). Last, a BVS data set was acquired using the same parameters and sequence (380 volumes; Figure 1A).

**Figure 1.**
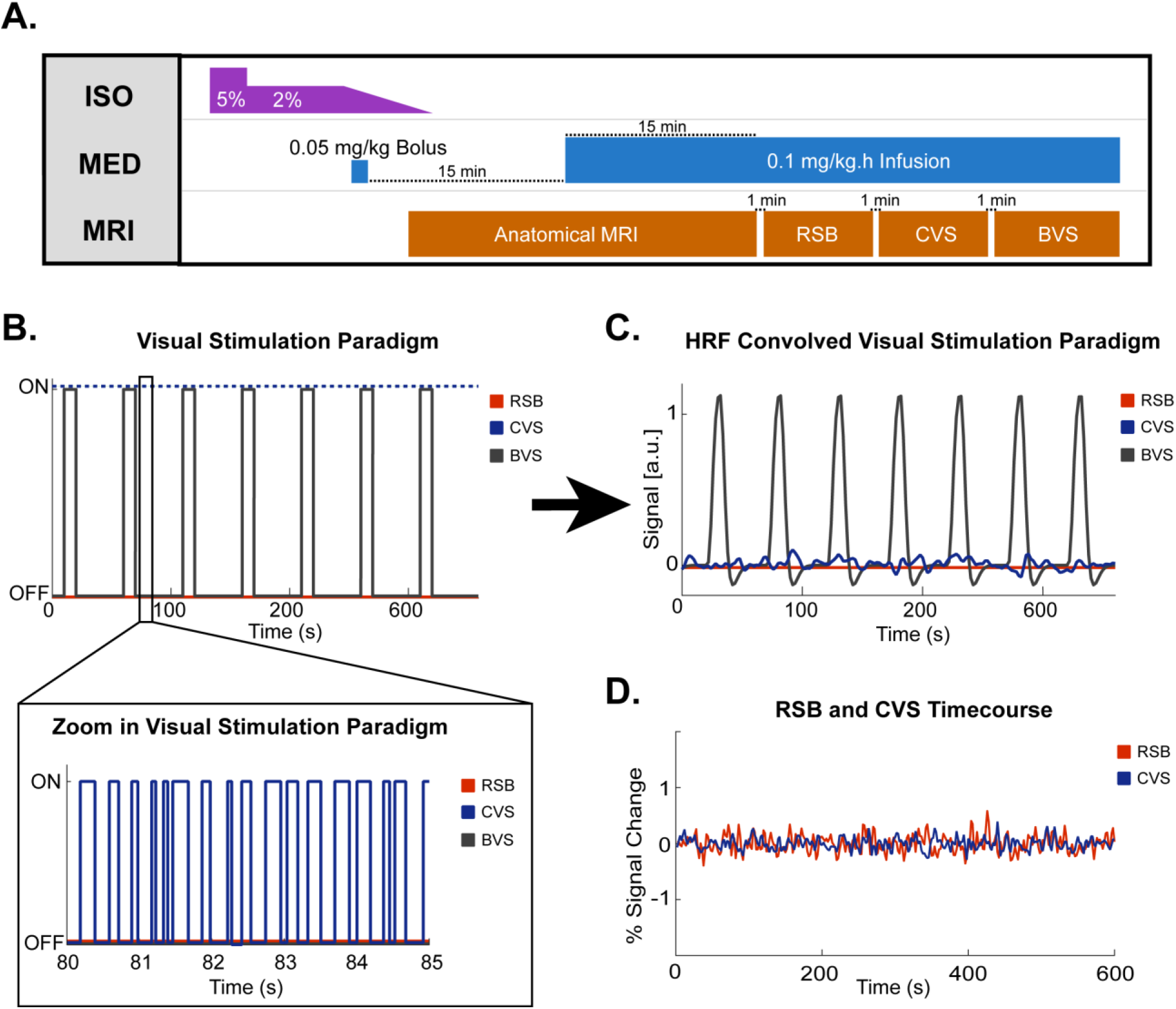
Scanning protocol and visual stimulation paradigms. A. Scanning protocol. For handling procedures, animals were first anesthetized using 5% isoflurane (ISO) for induction followed by 2% ISO for maintenance. Once the animal is fixated in the scanner bed, a bolus of 0.05mg/kg of medetomidine (MED) was administered and ISO anesthesia was gradually decreased to 0% ISO. After 15 minutes post bolus injection, a continuous infusion of 0.1 mg/kg.h MED was administered to the animal. For imaging procedures, first a set of anatomical Turbo RARE T2 scans were acquired and shimming procedures were performed. Next, 30 min post bolus injection a resting state baseline (RSB) scan was acquired. Subsequently, continuous visual stimulation (CVS) paradigm was turned on and after one minute the CVS scan was acquired. Lastly, after a recovery time of 1 minute a block design visual stimulation (BVS) fMRI scan was acquired. B. Visual stimulation paradigm of RSB, CVS and BVS scan indicating when visual stimuli was turned on or off. C. Haemodynamic Response Function (HRF) convolved visual stimulation paradigm of RSB, CVS and BVS scan showing predicted BOLD signal response in arbitrary units (a.u.) from each condition. D. Example of acquired RSB and CVS normalized BOLD signal time course in the visual network.

### 2.4. Visual stimulation

Visual stimulation with flickering light was presented using a white LED coupled to a fiber-optic cable, which was centrally placed in front of the animal’s head. The LED light was controlled by a voltage-gated device to control the triggering of the LED light (ON-OFF) driven by a RZ2 BioAmp Processor (Tucker-Davis, Alachua). Stimulus timing and alignment to the MR imaging was achieved by TTL pulses sent by the scanner at the beginning of every volume of the fMRI scan.

#### 2.4.1. Continuous visual stimulation

The CVS was used to induce a continuous visual sensory drive with randomization of light pulses to avoid sensory adaptation effects and to constantly trigger bottom-up attention mechanisms. This condition can be assumed to create a steady brain state similar to rest and thus could be analyzed the same way as the RSB scan (Figure 1B-D). The CVS was initiated 1 min before the start of the acquisition to avoid the detection of the initial transient increase of brain activity, which occurs when the visual stimulation is turned on. The stimulation paradigm was controlled via Matlab code (MATLAB R2014a, The MathWorks Inc. Natick, MA, USA) using an USB to serial port connection (IC-232A, Rotronic) and consisted of continuous short pulses of light and inter-light intervals (both with a random duration between 50-250 ms) (Figure 1B). Convolution of the CVS stimulus paradigm with the canonical hemodynamic response function (HRF) in SPM12 was performed in order to demonstrate that the expected signal fluctuations from CVS are negligible in comparison to those expected by the BVS paradigm (Figure 1C).

#### 2.4.2. Block design visual stimulation

To evoke BOLD responses from the visual system, fMRI scan was acquired during a block design paradigm with a visual stimulation frequency of 4 Hz, duty cycle 50% (125 ms ON/OFF), an initial OFF block of 10 seconds followed by 15 ON/OFF blocks of 10 s and 40 s respectively (Figure 1B-C).

### 2.5. Breathing rate processing

Breathing rate pressure signals were analyzed to investigate the potential influence of visual stimulation on the animals’ physiology. First, breathing rates were calculated for each volume by calculating the median period of the breathing cycle between the initial and the following volume and inverting this value to breaths per minute. For resting state data, averaged breathing rate from the complete scans were compared between the RSB and CVS condition using a paired t-test (p<0.05). For the BVS scans, averaged breathing rate over all visual stimulation blocks was compared ten seconds before with ten seconds during visual stimuli using a paired t-test.

### 2.6. MRI Processing

All data processing was performed using SPM 12 software (Statistical Parametric Mapping, http://www.fil.ion.ucl.ac.uk), REST toolbox (REST1.7, http://resting-fmri.sourceforge.net) and GIFT toolbox (Group ICA of fMRI toolbox version 3.0a: http://icatb.sourceforge.net/). Pre-processing consisted of realignment of the data towards the first image of each scan using a 6-parameter (rigid body) spatial transformation, normalization towards a study specific EPI template using a global 12-parameter affine transformation, followed by a non-linear transformation. Finally, data were smoothed in-plane using a Gaussian kernel with a full width at half maximum of twice the voxel size (0.458 × 0.626 mm). rsfMRI data were then further band pass filtered between 0.01-0.2 Hz using REST toolbox.

#### 2.6.1. Functional connectivity

##### 2.6.1.1. ICA components

To obtain the RSNs from the rsfMRI data, an independent component analysis (ICA) was performed on RSB data. First, movement was regressed out of the data based on the estimators from the realignment procedure using the REST toolbox. Next, ICA was performed in GIFT using the Infomax algorithm with predefined number of 15 components as used previously (Jonckers et al., 2011). Components representing functional networks were identified by comparison to previous observed RSNs and discarding a minority of artefactual components (e.g. only edge of the brain) by careful visual inspection (Gozzi and Schwarz, 2016; Jonckers et al., 2011; Lu et al., 2012; C.P. Pawela et al., 2008; Sierakowiak et al., 2015). Afterwards, the selected mean z-scored RSNs group components were thresholded at z-score > 1 to obtain a mask of each network.

##### 2.6.1.2. ICA-based inter-network connectivity

To evaluate inter-network connectivity, network masks were used as regions of interest (ROI) in correlation-based analysis per subject. Pairwise Pearson correlation coefficients between the average BOLD time series of each network ROI were calculated and Fisher z-transformed using an in-house Matlab program. Mean Fisher z-transformed inter-network correlation matrices were calculated for RSB and CVS conditions. Statistical analysis between conditions was performed using a repeated measures 2-way ANOVA (p<0.05, Sidak correction for multiple comparisons). To calculate the FC between the DMLN and visual network, z-transformed correlation values of the cingulate-retrosplenial, hippocampal and temporal-prefrontal with visual network were averaged. Differences in correlation between the two conditions were then investigated using pairwise t-test (p<0.05).

##### 2.6.1.3. ROI-based intra- and inter-network connectivity

To investigate within and between network connectivity differences, specific anatomically defined bilateral ROIs were selected in the DMLN (based upon the RSB ICA networks: cingulate cortex (Cg), retrosplenial cortex (RS), parietal association cortex (PtA), temporal association cortex (TeA)) as well as in the visual system (based upon the visual stimulation correlated ICA component of the BVS: lateral geniculate nucleus (LGN), superior colliculus (SC), visual cortex (VC)) (suppl. figure 2). Furthermore, a ROI in the primary somatosensory cortex (SS) was added as a control region. Pairwise Pearson correlation coefficients between the average BOLD time series of each ROI were calculated and Fisher z-transformed. Mean z-transformed ROI connectivity matrices were calculated for RSB and CVS conditions. To compare RSB and CVS condition, statistical analysis between conditions was performed using a repeated measures 2-way ANOVA (p<0.05, Sidak correction for multiple comparisons).

##### 2.6.1.4. Seed-based analysis

To investigate how the full-brain FC of the Cg is influenced by CVS, seed-based analysis was performed. To this end, the mean time course of Cg was used as a predictor in a General linear model (GLM) and motion parameters were included as covariates. Each subject’s FC between Cg and DMLN or visual network was extracted by using the respective binarized masks derived from the ICA and calculating the mean T-values within this mask for each individual subject. The DMLN network mask was constructed by the union of the cingulate-retrosplenial network mask, hippocampal network mask and temporal-prefrontal network mask. Statistical analysis to compare differences between the conditions was performed by means of a paired t-test (p<0.05).

#### 2.6.2. Block design fMRI

BVS data were processed using a group ICA approach using the GIFT toolbox with an Infomax algorithm and 15 predefined components. Group ICA analysis was chosen instead of the standard GLM to increase sensitivity towards detecting responding brain regions by not relying on predetermined BOLD HRF functions (Calhoun et al., 2009). The components temporal correlation with the visual stimulation paradigm was then calculated using R-square statistic in the GIFT toolbox. The component with highest temporal correlation with the visual stimulation paradigm were regarded as responding regions. Event related BOLD responses were extracted from a binary mask of responding regions, which are either correlated (z-score > 1) or anti-correlated (z-score < −1) with the ICA time course. To confirm our results, GLM analysis was performed for each subject within the ICA component mask (z-score > 1 and < −1) of the responding regions using a canonical HRF function in SPM 12. The model was set to either detect a positive BOLD response during the stimulation period, to locate visual responding regions, or during the rest period, to detect regions which are more active during rest vs stimulation period i.e. DMLN activity. One sample t-test (p<0.001, uncorrected for multiple comparisons) was performed to evaluate BSV group response.

## 3. Results

### 3.1. Block-design visual stimulation

To investigate whether activity in the DMLN of anesthetized rats is changing between visual sensory stimulation and rest periods, we analyzed the block-design fMRI experiment. To this end, we performed group-ICA analysis and sorted the components based on their temporal correlation with the visual stimulation paradigm. The component with the highest correlation (R^2^=0.64) showed clear BOLD signal increases during the ON-blocks indicating a strong visual activation of multiple responding regions (Figure 3A). The mean group statistical ICA map (z-score > 1 and < −1) of this component revealed that the activated regions were, as expected, areas involved in visual processing, *i.e.* the LGN, SC, VC (Figure 3B). Event related analysis of all regions which are correlated with the ICA time course demonstrated that visual stimulation evoked a positive BOLD response as expected (Figure 3C). Interestingly, we also observed regions such as the temporal association cortex/auditory cortex (TeA/AC) and RS, being anti-correlated to the ICA component’s time course, indicating higher BOLD signal during rest vs stimulation (Figure 3B). Event related analysis of anti-correlated regions with the ICA time course showed a subtle BOLD signal decrease during visual stimulation which rebounded and increased during blanks (Figure 3C). To confirm our results, GLM modeling was applied and detected increased BOLD response during visual stimulation in LGN, SC and VC as well as increased BOLD response during blanks in TeA/AC (suppl. figure 3).

**Figure 3.**
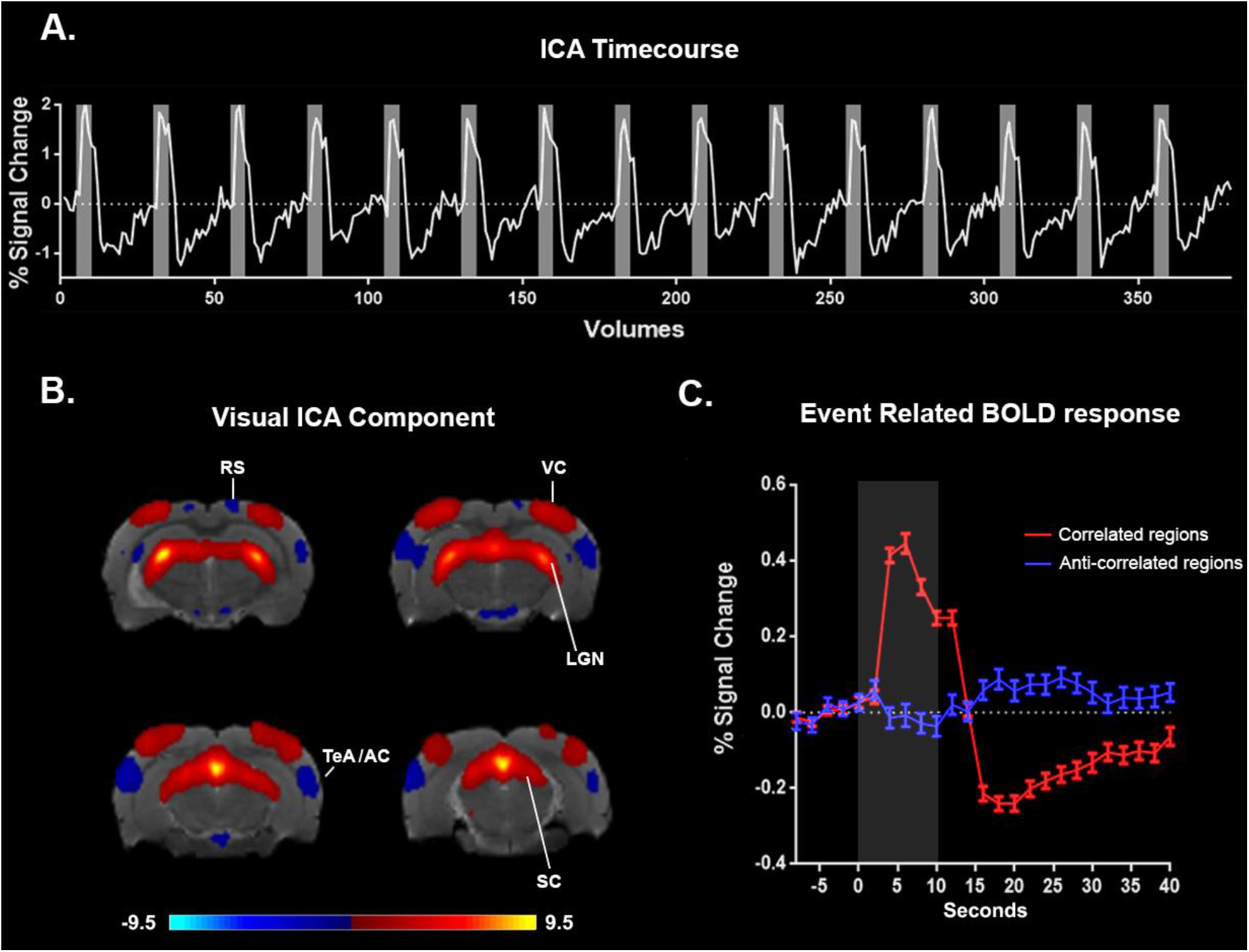
Functional MRI ICA. A. ICA Time course. Time course from mean group visual ICA component with highest temporal correlation (R^2^=0.64) with the visual stimulation paradigm (grey blocks). B. Visual ICA Component. Mean statistical group ICA map (z-score −1< and >1) demonstrating areas which are correlated to the mean ICA component’s time course involved in visual processing, i.e. visual cortex (VC), lateral geniculate nucleus (LGN) and superior colliculus (SC). Regions demonstrating a BOLD time course that is anti-correlated with the mean ICA component’s time course, meaning higher BOLD signal during rest than during visual stimulation, were observed in DMLN regions i.e. temporal association cortex/auditory cortex (TeA/AC) and retrosplenial cortex (RS). C. Event related response of the regions correlating (Red) and anti-correlating (Blue) to the ICA component’s time course. Grey block indicates visual stimulation period

### 3.2. Resting state vs Continuous Visual Stimulation

#### 3.2.1 Decreased intra- and inter-network connectivity during CVS

To identify potential changes in FC induced by our randomized continuous stimulation paradigm, we first performed group-ICA for the RSB condition and identified commonly observed RSNs. Functionally relevant components included three subnetworks of the DMLN: a cingulate-retrosplenial network (with cingulate cortex, retrosplenial cortex and parietal association cortex), a hippocampal network (with subiculum, dentate gyrus, CA1 and CA3), and a temporal-prefrontal network (with orbito-frontal cortex, prelimbic cortex and temporal association cortex). Furthermore, ICA analysis detected a visual network, a caudate putamen network, a primary and secondary somatosensory network, a barrel field network and a limbic network (including the amygdala and ventral striatum). Next, to evaluate intra- and inter-network connectivity, we threshold each ICA-component map (z-values > 1), extracted the time-courses and performed pairwise correlations for both the RSB and CVS conditions (Figure 5). Statistical analysis comparing RSB and CVS condition using a repeated measures 2-way ANOVA (multiple comparison correction Sidak p<0.05) detected a significantly decreased intra-DMLN network correlation in the CVS condition (*i.e* subnetworks cingulate-retrosplenial and temporal-prefrontal (p<0.001)) as well as decreased inter-network correlations between DMLN subnetworks and the visual network (*i.e.* cingulate-retrosplenial network with visual network (p=0.002) and temporal-prefrontal network with visual network (p=0.027).

**Figure 4.**
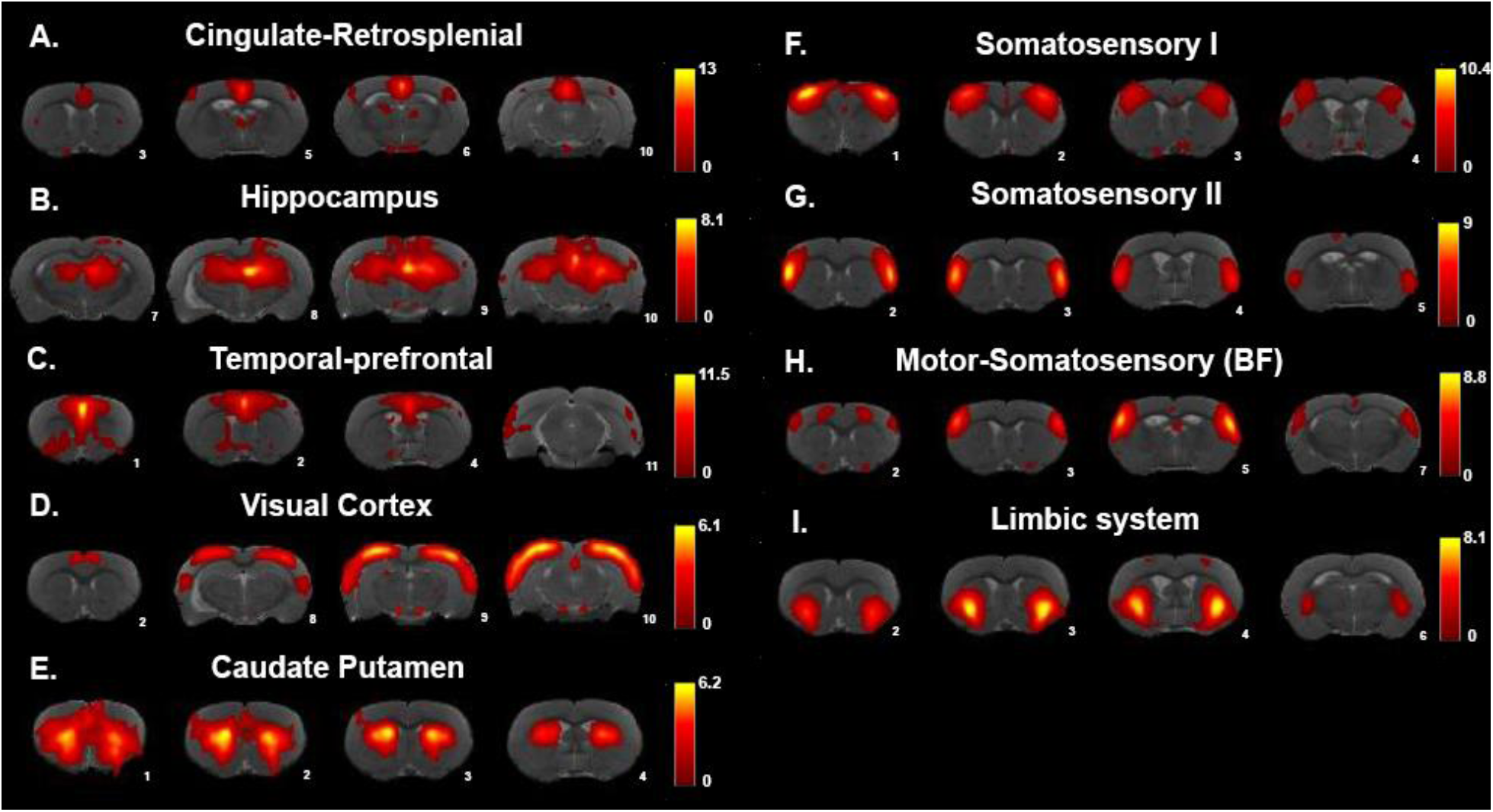
ICA of Resting State Baseline (RSB) condition. Mean group statistical ICA maps (z-score>1) revealed nine functionally relevant components. A.-C. The DMLN in rats split up in three subcomponents i.e. A. Cingulate-Retrosplenial Network (with cingulate, retrosplenial cortex and parietal association cortex), B. Hippocampal network (with subiculum, dentate gyrus, CA1 and CA3) and C. Temporal-prefrontal network (with orbito-frontal cortex,prelimbic cortex and temporal association cortex). D. Visual network (with visual and somatosensory cortex). E. Caudate Putamen network. F.-G. Primary and secondary somatosensory network. H. Barrel field network. I. Limbic network (with amygdala and ventral striatum).

**Figure 5.**
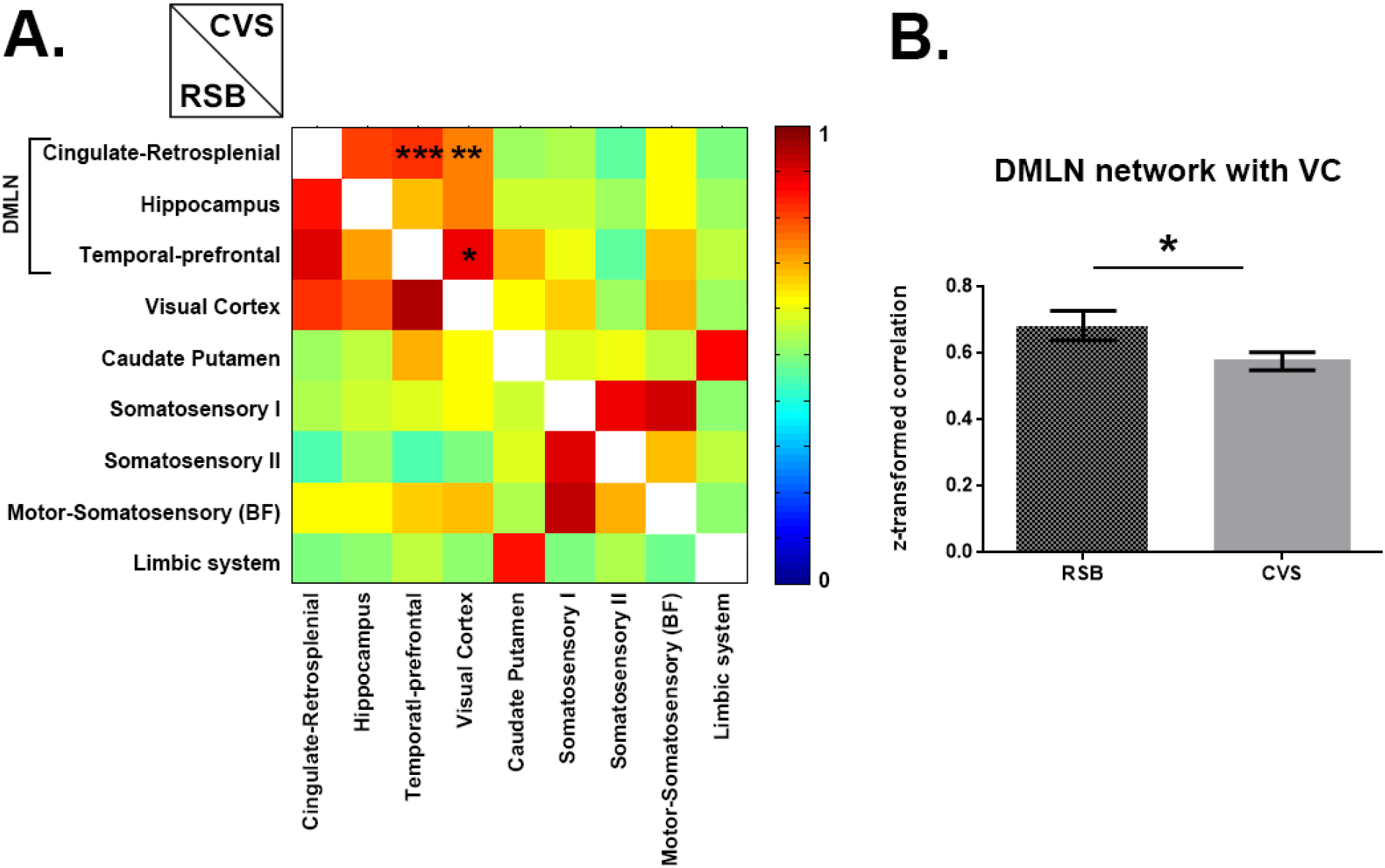
Inter-network functional connectivity. A. Pairwise z-transformed Pearson correlation matrix of network components’ time courses of the resting state baseline (RSB) (Bottom) and of the continuous visual stimulation (CVS) condition (Top). Stars indicate significant differences found between the two conditions with repeated measures 2-way ANOVA (p<0.05, with Sidak multiple comparison correction). For CVS, significant decreased inter-network connectivity was found between subnetworks of the DMLN (i.e cingulate-retrosplenial network and temporal-prefrontal network) and between DMLN subnetworks and visual network. Color bar represents z-values. *p<0.05, **p<0.005, ***p<0.001 B. Inter-network connectivity between DMLN and Visual network (average of connectivity between cingulate-retrosplenial network and visual network, hippocampal network and visual network, and temporal-prefrontal network and visual network). Statistical analysis using a paired t-test detected a significant decreased inter-network connectivity in the CVS condition as compared to the RSB condition (p< 0.05).

Since CVS induced decreased inter-network correlation between the DMLN subnetworks and the visual network, we further assessed the pairwise correlation between specific ROIs of these networks to zoom-in and better understand the sources of these decreases. ROIs in DMLN (*i.e.* Cg, RS, TeA and PtA) and visual system (*i.e.* LGN, SC and VC) were selected. In addition, we included the SS as a control area. Pairwise correlation between each ROI’s averaged BOLD time course was performed and compared between RSB and CVS condition (Figure 6). Statistical analysis using a repeated measures 2-way ANOVA (Sidak multiple comparisons correction p<0.05) showed a decreased correlation between the DMLN ROIs *i.e.* Cg-RS (p<0.001), Cg-PtA (p=0.005), RS-TeA (p=0.002) and RS-PtA (p=0.006) as well as between DMLN and visual ROIs *i.e.* Cg-VC (p<0.001), RS-VC (p=0.005), PtA-SC (p=0.036), and PtA-VC (p=0.002). None of these areas showed a significant change in correlation with the SS.

**Figure 6.**
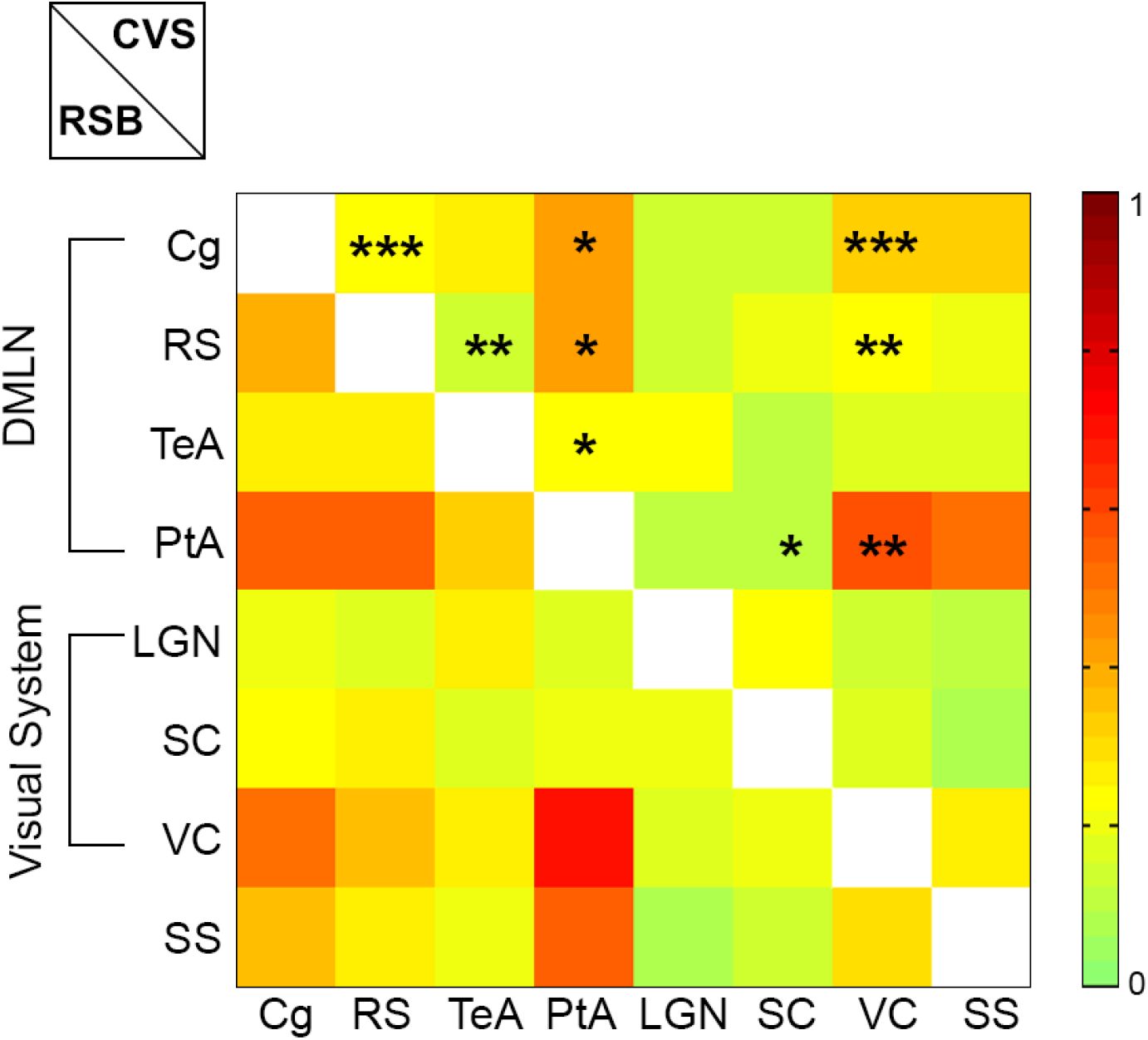
ROI based analysis. A. Pairwise z-scored Pearson functional connectivity (FC) matrix between time courses of ROIs of the DMLN (i.e. cingulate cortex (Cg), retrosplenial cortex (RS), temporal association cortex (TeA) and parietal association cortex (PtA)), the visual system (i.e. lateral geniculate nucleus (LGN), superior colliculus (SC) and visual cortex (VC)) and the somatosensory cortex (SS) as a control region. Top half of the matrix represents FC of continuous visual stimulation (CVS) condition. Bottom half of the matrix represents FC of resting state baseline (RSB) condition. Color bar represents z-values. Stars indicate significant differences found between the two conditions (diagonally symmetric positions) with repeated measures 2-way ANOVA (p<0.05, with Sidak multiple comparison correction). Decreased FC was detected between DMLN ROIs i.e. Cg-RS, Cg-PtA, RS-TeA and RS-PtA as well as between DMLN and visual system ROIs i.e. Cg-VC, RS-VC, Pta-SC and PtA-VC. *p<0.05, **p<0.005, ***p<0.001

#### 3.2.2 Voxel-based functional connectivity of Cingulate cortex

The cingulate cortex, a major node of both DMLN in rodents as well as DMN in humans has been shown to change its activity during unpredictable stimuli (Hahn et al., 2007). Since our stimuli in the CVS condition were similarly designed to be unpredictable we selected this area for seed-based analysis. As demonstrated in the statistical FC maps (one sample t-test, p<0.001, family wise error (FWE) corrected for multiple comparisons) presented in Figure 7A, the Cg demonstrated wider brain connectivity during the RSB in comparison to the CVS condition. To further quantify this effect, we performed a paired t-test for the T-values within the DMLN or visual network. We found that CVS condition induced a decreased correlation between Cg and DMLN as well as between Cg and the visual network (Figure 7B-C).

**Figure 7.**
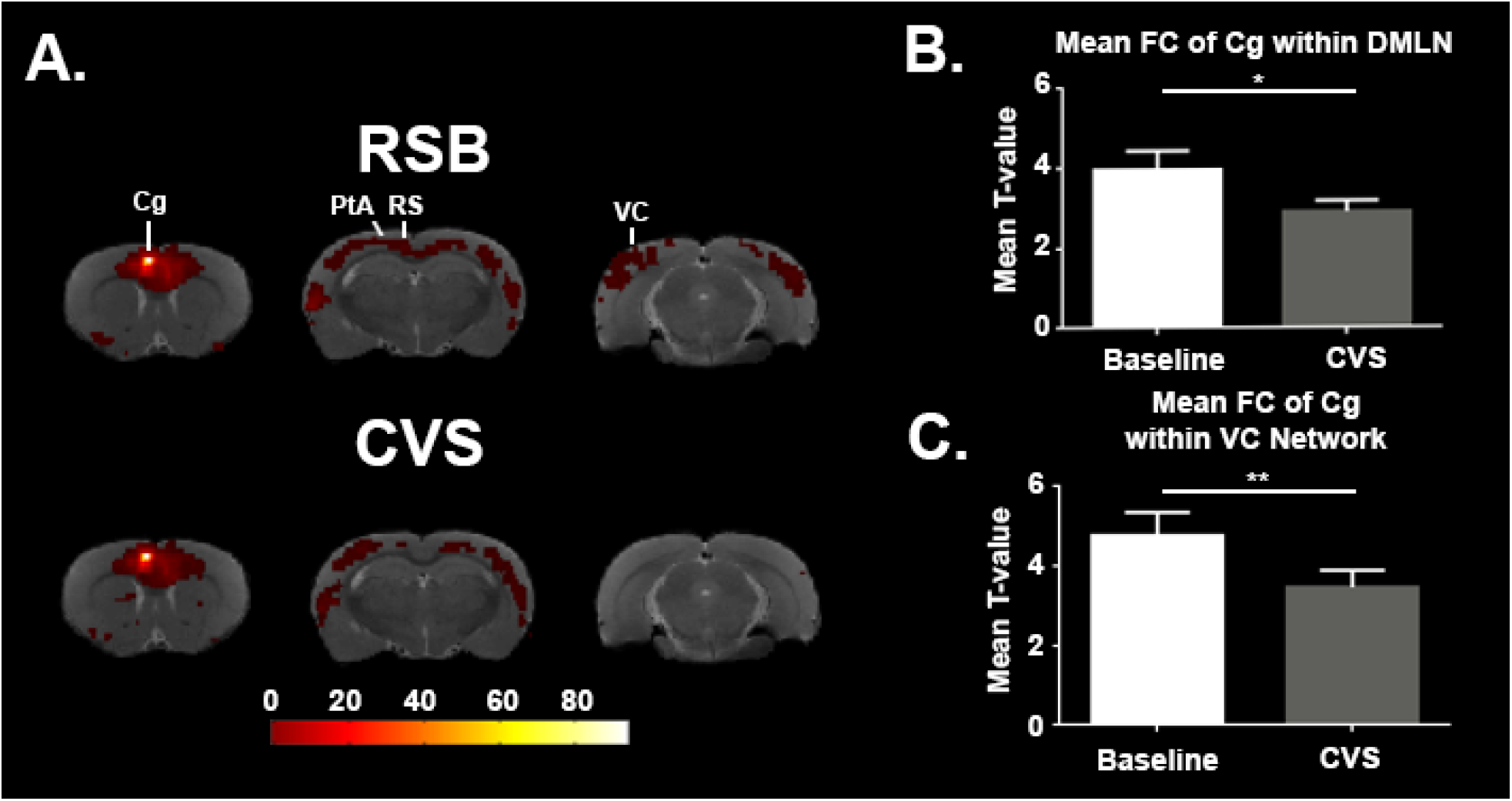
Seed based analysis of functional connectivity with the cingulate cortex as seed region. A. Statistical maps of functional connectivity (p<0.05 with family wise error correction (FWE) for multiple comparison correction) of the cingulate cortex (Cg) in the resting state baseline (RSB) condition (Top) and continuous visual stimulation (CVS) condition (Bottom). Decreased FC in the parietal association cortex (PtA), retrosplenial cortex (RS- and visual cortex (VC) was observed in the CVS condition. Color bar represents t-values. High t-values indicate high functional connectivity with the seed B. Bar graph of the mean T-value with ± SEM within the default mode like network. For each subject, mean T-values were extracted within the default mode like network ICA mask. Results show that CVS significantly decreased connectivity of the Cg (paired t-test, p<0.05) with the default mode like network. C. Bar graph of the Mean T-value with ± SEM of all subjects within the Visual network (VC). For each subject, mean T-values were extracted within the VC network ICA mask. Results show that continuous visual stimulation (CVS) significantly decreased connectivity of the Cg (paired t-test, p<0.05) with the VC network.

### 3.3 Unaltered breathing rate during visual stimulation

The pressure signal for breathing rate of the animals was recorded throughout the whole experiment and analyzed to assess potential changes on the general physiological state due to the visual stimulation. A paired t-test showed that there were no significant changes in breathing rate due to the visual stimulation (p=0.79) (suppl. figure 4A). Breathing rate was further compared between RSB and CVS conditions using a paired t-test (p<0.05). Likewise, CVS did not significantly alter the breathing rate (p=0.1) (suppl. figure 4B).

## 4. Discussion

In the current study, we investigated the impact of visual stimulation on the DMLN activity and its FC in rats. We found that visual stimulation could deactivate nodes of the DMLN and could decrease FC within DMLN as well as across DMLN and visual networks.

### 4.1. Block design visual stimulation induces deactivation in DMLN regions

Block design visual stimulation was performed a) to identify the visually responsive areas and b) to investigate the influence of visual stimulation on the DMLN.

Similar to previous visual fMRI studies in rats, we detected activation of visual processing areas including LGN, SC, and VC (Van Camp et al., 2006), (Chrisopher P. Pawela et al., 2008). In addition to these activating regions, we detected deactivating regions that demonstrated reduced activity during visual stimulation and displayed increased activity during rest i.e. TeA/AC and RC. Interestingly, these areas have been shown to be nodes of the DMLN in rats (Lu et al., 2012). This finding seems in contrast to earlier human studies that did not detect significant reductions in DMN activity during passive sensory processing states that have low cognitive demand e.g. flashing checkerboard pattern presentation (Greicius et al., 2003). However, a number of important differences in our study could explain this discrepancy. Firstly, it should be noted that the effect we found in rats was very subtle. Thus, it is possible that similar effects could be present in humans but not sufficiently strong to be detected during awake conditions that are potentially compromised by additional processes. Previous studies directly demonstrated that the magnitude and extent of the suppression depends on the difficulty of the cognitive task (Leech et al., 2011; Mayer et al., 2010). As our study was performed in anesthetized rats, although not optimal for top-down cognitive processing, could be beneficial for identifying subtle bottom-up effects that could be otherwise hindered by additional variability induced by awake conditions. An alternative explanation is that in rodents passive sensory stimulation is more cognitively demanding in comparison to humans. Visual stimulation in rodents is thought to increase behavior mechanisms such as fear, which could be responsible for modulating the DMLN (McClearn, 1960). As previous studies in human have shown, fear can readily deactivate the DMN (Marstaller et al., 2017). Our observation could therefore suggest stronger cognitive involvement in rats during visual stimulation (Anticevic et al., 2013).

### 4.2. Continuous visual stimulation decreases inter- and intra-network FC of the DMLN

We explored the reorganization of functional networks during a visually stimulated brain state induced by CVS. The CVS paradigm was specifically developed to get the animal in a visually attentive stimulated steady state and stochasticity was included to avoid habituation towards the stimulus. Firstly, ICA analysis was performed on the RSB data in order to identify the RSNs. These networks showed strong bilateral connectivity and were similar to previously described networks in rats (Hutchison et al., 2010; Jonckers et al., 2011). We then investigated how the activity and connectivity within and across these networks changed in the CVS condition.

By comparing FC between networks in both conditions, we demonstrated that CVS decreased inter-network FC between DMLN subcomponents (cingulate-retrosplenial and temporal-prefrontal networks (Lu et al., 2012)) and with the visual network. The observed decrease in inter-network connectivity during the CVS condition suggests an alteration in communication between the DMLN and the visual network. This finding could subserve the enhancement of local, input specific visual processing during CVS versus a higher inter-network communication during rest conditions.

Further, we performed ROI based analysis focusing on areas within the DMLN. This analysis demonstrated that intra-DMLN FC was also decreased. This included connections between multiple major nodes of the DMLN *i.e* Cg-RS, Cg-PtA, RS-PtA and RS-TeA. Similar to activity level decreases, connectivity decreases within human DMN were previously observed only with tasks involving higher cognitive load (Elton and Gao, 2015; Fransson, 2006; Gao et al., 2013; Marrelec and Fransson, 2011), while simple visual stimulation with a constantly flickering checkerboard pattern were not able to induce such deactivation (Di et al., 2015). Interestingly, the results observed in our study are more consistent with human data from subjects in a higher attentive and cognitive brain state.

## 5. Conclusion

In summary, we demonstrated that simple yet stochastic sensory stimulation in anesthetized rats could a) deactivate certain nodes of the DMLN, and b) reduce intra- and inter-DMLN network connectivity simulating similar results in humans performing task involving high cognitive and top-down attentional demand. We conjecture that the stochasticity of our stimulus, may play an important role in consistently and continuously driving bottom-up attention triggering mechanisms. Given that the bottom-up and top-down attentional systems share specific network components (Katsuki and Constantinidis, 2014), we suggest that both attention mechanisms are able to deactivate and reduce functional connectivity of the DMN. These results are very significant and could prove immensely useful not only for our better understanding of the DMN using animal models but are also very promising for being used in human patients that are anesthetized or non-responsive as a result of trauma and or injury. However, to more explicitly demonstrate the link between attention systems and DMN activity and connectivity, more studies are required both in humans using stimuli specifically designed to continuously drive bottom-up attention, as well as in awake and behaving animals including more complicated cognitive tasks.

## Supporting information

## Acknowledgements

This research was supported by the fund of scientific research Flanders (FWO G048917N), Flagship ERA-NET (FLAG-ERA) FUSIMICE (grant agreement G.0D7651N) and University Research Fund of University of Antwerp (BOF DOCPRO FFB150340).

## Abbreviations

AC: Auditory cortex
BOLD: Blood oxygen level dependent
BVS: Blocked visual stimulation
Cg: Cingulate cortex
CVS: Continuous visual stimulation
DMLN: Default mode like-network
DMN: Default mode network
FC: Functional connectivity
fMRI: Functional magnetic resonance imaging
FOV: Field of view
FWE: Family wise error
GLM: General linear model
HRF: Hemodynamic response function
ICA: Independent component analysis
ISO: Isoflurane
LGN: Lateral geniculate nucleus
MD: Matrix dimensions
MED: Medetomidine
PtA: Parietal association cortex
ROI: Regions of interest
RS: Retrosplenial cortex
RSB: Resting state baseline scan
rsfMRI: Resting-state functional magnetic resonance imaging
RSNs: Resting state networks
SC: Superior colliculus
SS: Somatosensory cortex
ST: Slice thickness
TE: Echo time
TR: Repetition time
VC: Visual cortex

