## Supplementary material for "Bottom-up sensory processing can decrease activity and functional connectivity in the default mode like network in rats"

### Supplementary figures

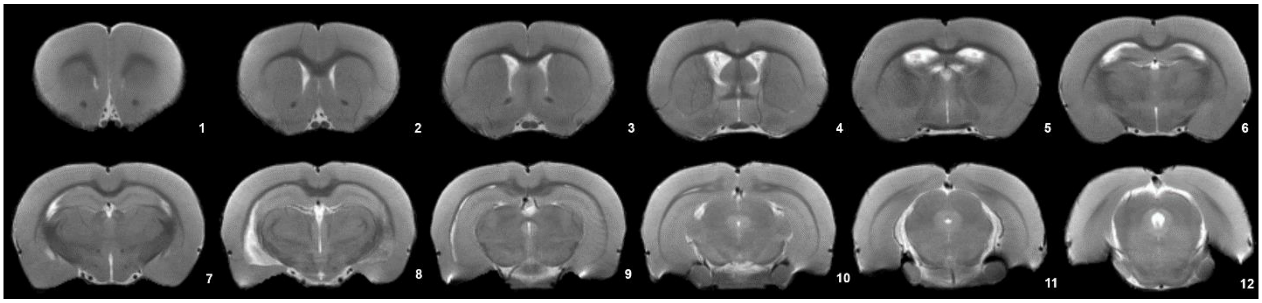


***Supplementary figure 1. Turbo RARE T2-weighted slice package.*** *Slices were selected to cover the brain from approximately 3.3 mm anterior to bregma to 7.5 mm posterior to bregma.*

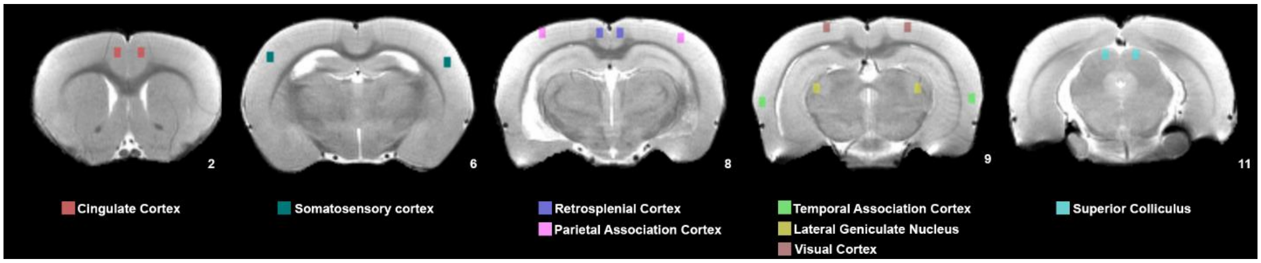


***Supplementary figure 2.*** *Region of Interest (ROI) selection. For ROI based analysis, left and right ROIs were selected within the DMLN i.e. cingulate cortex (red), retrosplenial cortex (purple), parietal association cortex (pink) and temporal association cortex (green) as well as ROIs within the visual system i.e. lateral geniculate nucleus (yellow), visual cortex (brown) and superior colliculus (cyan).*

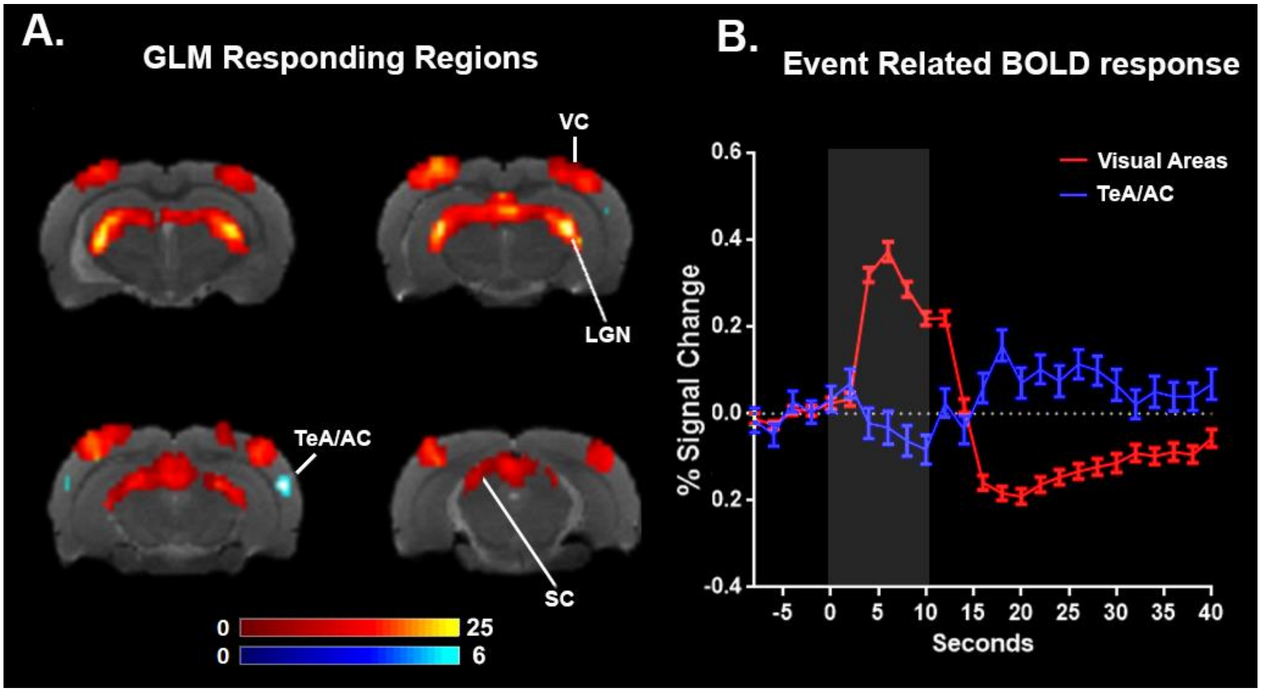


***Supplementary figure 3. Functional MRI GLM*** *A. Mean statistical T-map resulting from the general linear modeling analysis. During visual stimulation (warm colors), positive BOLD response was detected in visual processing areas i.e. visual cortex (VC), lateral geniculate nucleus (LGN) and superior colliculus (SC).During rest (cold colors), regions of DMN i.e. temporal association cortex/auditory cortex (Tea/AC) demonstrated higher activity than during visual stimulation. C. Event related response of the visual areas (Red) and Tea/AC (Blue). Grey block indicates visual stimulation period.*

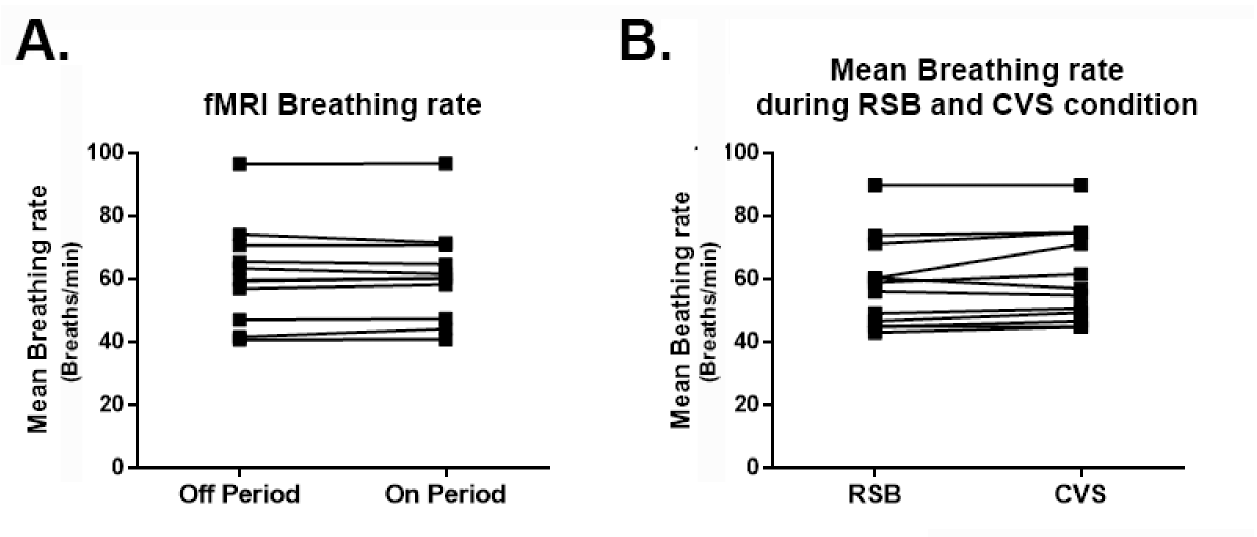


***Supplementary figure 4.*** *Breathing rate during visual stimulation. A. Breathing rate during block design visual stimulation. Breathing rate was analyzed by comparing the averaged breathing rate 10s before visual stimulation with the average breathing rate during the 10s of all blocks of the visual stimulation. No significant difference in breathing rate was observed due to visual stimulation (*$p>0.05$*). B. Breathing rate during resting state baseline (RSB) and continuous visual stimulation (CVS). Average breathing rate was compared for RSB and CVS condition. No significant difference in breathing rate was observed due to CVS (*$p>0.05$*).*
